# Omicron booster in ancestral strain vaccinated mice augments protective immunities against both the Delta and Omicron variants

**DOI:** 10.1101/2022.02.19.481110

**Authors:** Liqiu Jia, Yang Zhou, Shaoshuai Li, Yifan Zhang, Dongmei Yan, Wanhai Wang, Wenhong Zhang, Yanmin Wan, Chao Qiu

## Abstract

A booster vaccination is called for constraining the evolving epidemic of SARS-CoV-2. However, the necessity of a new COVID-19 vaccine is currently unclear. To compare the effect of an Omicron-matched S DNA vaccine and an ancestral S DNA vaccine in boosting cross-reactive immunities, we firstly immunized mice with two-dose of a DNA vaccine encoding the spike protein of the ancestral Wuhan strain. Then the mice were boosted with DNA vaccines encoding spike proteins of either the Wuhan strain or the Omicron variant. Specific antibody and T cell responses were measured at 4 weeks post boost. Our data showed that the Omicron-matched vaccine efficiently boosted RBD binding antibody and neutralizing antibody responses against both the Delta and the Omicron variants. Of note, antibody responses against the Omicron variant elicited by the Omicron-matched vaccine were much stronger than those induced by the ancestral S DNA vaccine. Meanwhile, CD8^+^ T cell responses against both the ancestral Wuhan strain and the Omicron strain also tended to be higher in mice boosted by the Omicron-matched vaccine than those in mice boosted with the ancestral S DNA vaccine, albeit no significant difference was observed. Our findings suggest that an Omicron-matched vaccine is preferred for boosting cross-reactive immunities.

## Introduction

The highly mutated SARS-CoV-2 Omicron (B.1.1.529) variant has been shown to substantially evade the neutralizing antibody responses elicited by current vaccines and early pandemic Alpha, Beta, Gamma, or Delta variant (1-9). A third dose of either homologous or heterologous COVID-19 vaccine was reported to enhance neutralizing antibody responses against the Omicron variant (3, 4, 10-15). However, the magnitudes of neutralizing activities towards Omicron after the booster dose were still far lower compared to earlier variants of concern (3, 5, 11, 16-19). Therefore, more effective vaccines or vaccination strategies are urgent in need to control the evolving pandemic of SARS-CoV-2 (20-22). Recently preprinted studies demonstrated that Omicron infection of previously vaccinated individuals could boost broadly neutralizing antibodies against different SARS-CoV-2 variants (23, 24), suggesting that the spike protein of the Omicron variant might serve as a good candidate antigen for a new COVID vaccine. Meanwhile, contradictory findings suggest that Omicron-matched vaccination shows no superiority in protection compared to immunization with current vaccines (25-28), which exaggerates concern about the effect of original antigenic sin caused by exposures to ancestral SARS-CoV-2 variants (29).

To investigate the effect of using Omicron-matched spike protein as a booster antigen, we firstly immunized mice with two-dose of a DNA vaccine encoding the spike protein of the ancestral Wuhan strain. Then the mice were boosted with DNA vaccines encoding spike proteins of either the Wuhan strain or the Omicron variant. Our data showed that the Omicron S DNA vaccine boosted cross-reactive antibody and T cell responses more efficiently than the ancestral S DNA booster vaccine.

## Materials and methods

### Ethical statement

All experiments and methods were performed in accordance with relevant guidelines and regulations. Mice experiments were reviewed and approved by the Research Ethics Review Committee of Shanghai Public Health Clinical Center.

### Constructions and preparation of candidate DNA vaccine encoding the full-length spike proteins of Wuhan or Omicron strain

The full-length *s* genes of the SARS-CoV-2 Wuhan and Omicron strain were optimized according to the preference of human codon usage and synthesized by Genewiz (Genewiz Biotech Co., Ltd., Suchow, China). The codon optimized spike genes were subcloned into the pJW4303 eukaryotic expression vector (Kindly gifted by Dr. shan Lu’s laboratory at the University of Massachusetts). The sequence of insertion was confirmed by Sanger sequencing (Sangon Biotech Co., Ltd., Shanghai, China). The recombinant plasmids for mouse vaccination were prepared using an EndoFree Plasmid Purification Kit (Cat# 12391, Qiagen, Hilden, USA).

### Mouse vaccination

Female C57BL/6J mice, 6-8-week-old, were purchased from Vital River Laboratory Animal Technology Co., Ltd. (Beijing, China) and housed in the SPF animal facility of Shanghai Public Health Clinical Center. The schedule of vaccination is shown in Figure 1. Briefly, 20μg of the S_Wuhan DNA vaccine was injected intramuscularly into each mouse at Week 0 and Week 2. Subsequently, the mice were boosted with 50μg of either the S_Omicron or the S_Wuhan DNA vaccine at Week 6. The control group was boosted with PBS. Four weeks after the final vaccination, the mice were euthanized. Peripheral blood and splenocytes were collected for assays of S protein specific immune responses.

**Figure 1.**
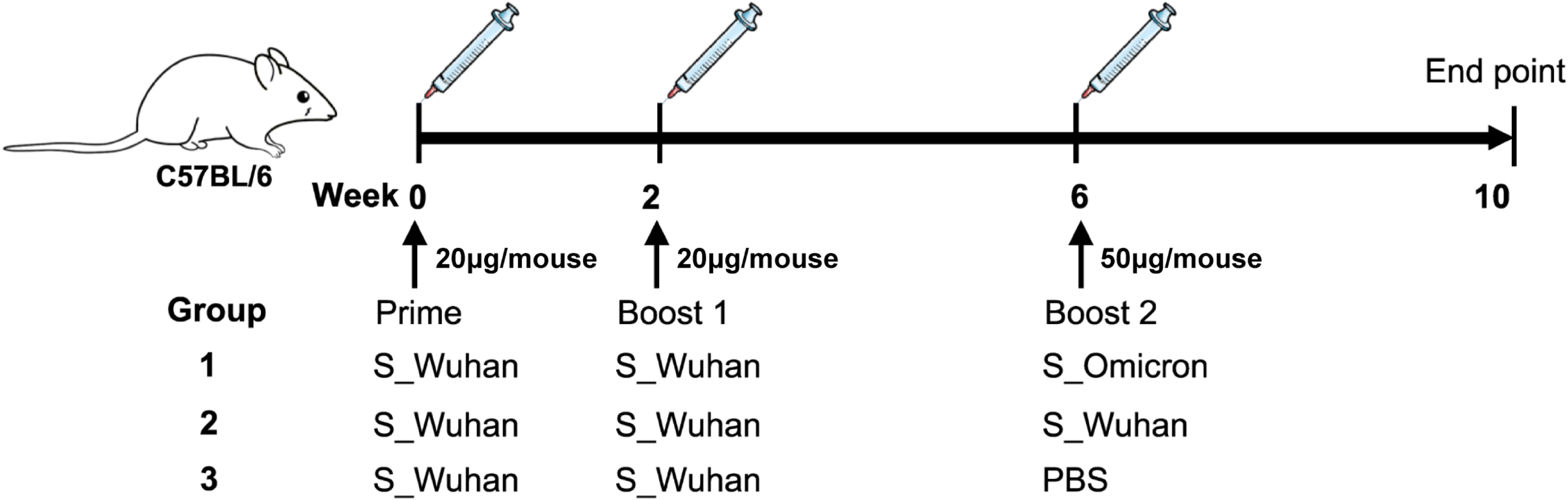
Schematic illustration of the vaccination schedule. Female C57BL/6 mice were injected intramuscularly with 20μg of the S_Wuhan DNA vaccine at Week 0 and Week 2. Subsequently, the mice were boosted with 50μg of either the S_Omicron (n=8) or the S_Wuhan (n=8) DNA vaccines at Week 6. The control group was boosted with PBS (n=6). Four weeks after the final vaccination, the mice were euthanized. Peripheral blood and splenocytes were collected on spot for assays of S protein specific immune responses.

### Detection of RBD specific binding antibodies

An in-house enzyme-linked immunosorbent assays (ELISA) were developed to measure RBD of Wuhan or Omicron strain of SARS-CoV-2 specific binding antibodies. High-binding 96-well EIA plates (Cat# 9018, Corning, USA) were coated with purified SARS-CoV-2 RBD protein of Wuhan strain (Cat# 40592-V08B, Sino Biological, China) or RBD protein of Omicron strain (Cat# 40592-V08H122, Sino Biological, China) at a final concentration of 1µg/ml in carbonate/bi-carbonate coating buffer (30mM NaHCO_3_,10mM Na_2_CO_3_, pH 9.6). After coating overnight at 4°C, the plates were blocked with 1×PBS containing 5% milk for 1 hour at 37°C. Subsequently, 100μl of serial dilutions of mouse serum was added to each well. After 1 hour incubation at 37°C, the plates were washed with 1×PBS containing 0.05% Tween 20 for 5 times. Then, 100μl of a HRP labeled goat anti-mouse IgG antibody (Cat# 115-035-003, Jackson Immuno Research, USA) diluted in 1×PBS containing 5% milk were added to each well and incubated for 1 hour at 37°C. After a second round of wash, 100μl of TMB substrate reagent (Cat# MG882, MESGEN, China) was added to each well. 6 minutes later, the color development was stopped by adding 100μl of 1M H_2_SO_4_ to each well and the values of optical density at OD_450nm_ and OD_630nm_ were measured using 800 TS microplate reader (Cat# 800TS, Biotek, USA).

### Flow cytometry assay

S specific T cell responses in splenocytes were detected by flowcytometry assay. Briefly, fresh splenocytes were prepared and stimulated with R10 containing synthesized peptides (0.66μg/ml for each peptide) covering the full-length spike protein of Wuhan strain or the fragments of Omicron spike protein containing mutations in round-bottom 96-well plates. 2 hours later, brefeldin A were added to each well at final concentrations of 1μg/ml. After 12 hours, the cells were collected and stained sequentially with Live/Dead dye (Fixable Viability Stain 510, Cat# 564406, BD Pharmingen) for 15 min at room temperature, surface markers (PE/Cyanine7-labeled anti-mouse CD3, Cat# 100220, BioLegend; APC-labeled anti-mouse CD4, Cat# 100412, BioLegend; PE-labeled anti-mouse CD8, Cat# 100708, BioLegend) for 30min at 4 °C and intracellular markers (BV421-labeled anti-mouse IFN-γ, Cat# 505830, BioLegend; FITC-labeled anti-mouse IL-2, Cat# 503806, BioLegend; BV711-labeled anti-mouse TNF-α, Cat# 506349, BioLegend) for 30min at 4 °C. After washing, the stained cells were resuspended in 200μl 1×PBS and analyzed using a BD LSRFortessa™ Flow Cytometer. The data were analyzed using the FlowJo 10 software (BD Biosciences, USA). The gating strategy was shown in Supplementary figure 1.

### SARS-CoV-2 pseudovirus neutralization assay

VSV-backboned SARS-CoV-2 pseudo-viruses were prepared according to a reported method (30). The neutralization assay was conducted by following the previously described procedure (30, 31). Briefly, 100μl of serially diluted mice sera were added into 96-well cell culture plates. Then, 50μl of pseudo-viruses with a titer of 13000 TCID_50_/ml were added into each well and the plates were incubated at 37°C for 1 hour. Next, Vero cells were added into each well (2×10^4^ cells/well) and the plates were incubated at 37°C in a humidified incubator with 5% CO_2_. 24 hours later, luminescence detection reagent (Bright-Glo™ Luciferase Assay System, Promega, USA) was added to each well following the manufacturer’s instruction. The luminescence was measured using a luminescence microplate reader (GloMax® Navigator Microplate Luminometer, Promega, USA) within 5 minutes. The Reed-Muench method was used to calculate the virus neutralization titer. Antibody neutralization titers were presented as 50% maximal inhibitory concentration (IC_50_).

### Statistical analysis

All statistical analyses were performed using GraphPad Prism 8 (GraphPad Software, Inc., La Jolla, CA, USA). Comparisons between two groups were conducted by the method of *t*-test. *P*<0.05 was considered as statistically significant.

## Results

### Boosting with a DNA vaccine encoding the spike protein of Omicron variant elicited cross-reactive antibody responses in mice

As aforementioned, Female C57BL/6J mice were immunized with 20μg of the S_Wuhan DNA vaccine at Week 0 and Week 2. Subsequently, the mice were boosted with 50μg of DNA vaccines expressing the S proteins of either the Omicron variant or the Wuhan strain (Figure 1) at Week 6. Sera were collected at the 4^th^ week post the last vaccination to measure RBD specific binding antibodies by a method of ELISA (Figure 2A and 2B). Our data showed that both the S_Wuhan and the S_Omicron DNA vaccines enhanced the RBD binding antibody responses compared with mice boosted with PBS. A booster shot of the S_Omicron DNA vaccine elicited stronger RBD binding antibody responses against the Omicron variant compared to a booster dose of the S_Wuhan DNA vaccine (P=0.004). The mean titer of RBD binding antibodies against the Wuhan strain had a tendency to be higher in the group boosted with the S_Omicron DNA vaccine than the group boosted with the S_Wuhan DNA vaccine, although the difference is not statistically significant. Then the neutralizing antibody titers were assessed using a VSV-backboned SARS-CoV-2 pseudo-virus assay (Figure 2C and 2D). Similar to the results of binding antibody assays, we found that a booster shot of the Omicron-matched DNA vaccine elicited stronger neutralizing antibody responses against the Omicron variant compared with booster doses of the S_Wuhan DNA vaccine and the PBS control (*P* =0.0047). The mean titer of neutralizing antibodies against the pseudo-virus of Delta also tended to be higher in the group boosted with the S_Omicron DNA vaccine than the group boosted with the S_Wuhan DNA vaccine, although the difference is not statistically significant.

**Figure 2.**
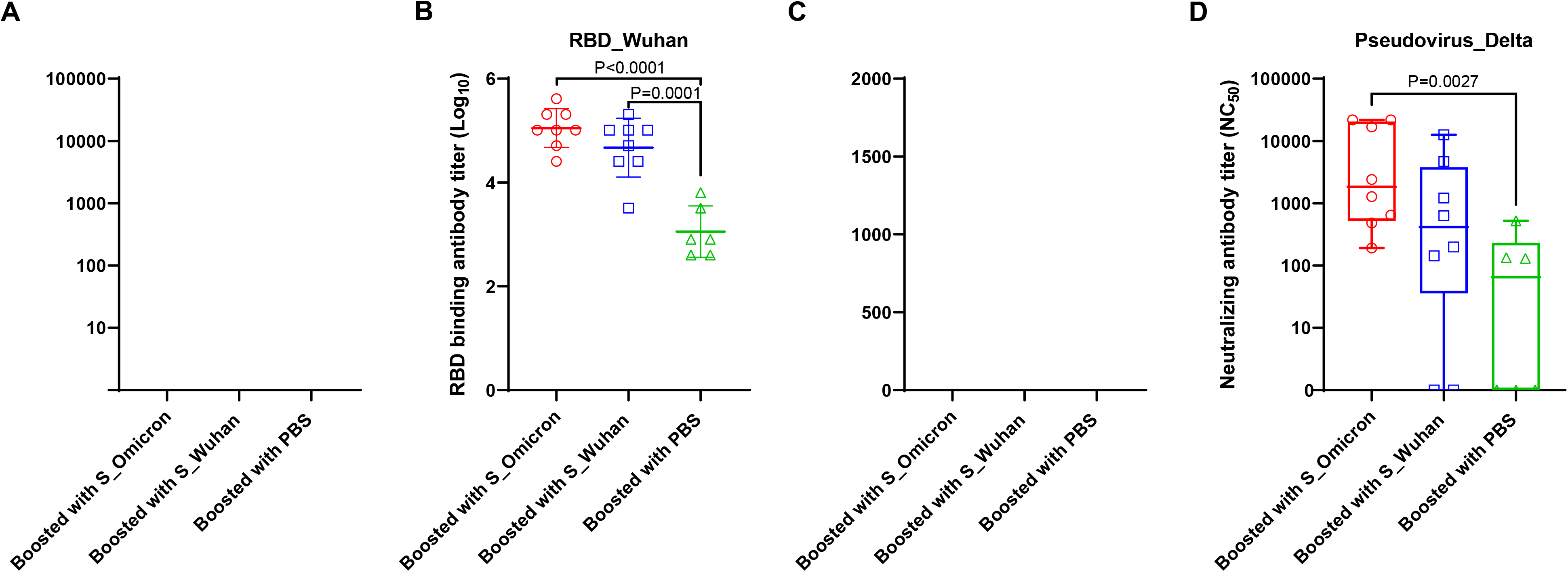
Boosting with the S_Omicron DNA vaccine elicited cross-reactive antibodies in mice. **A**, Titers of IgG binding to the RBD protein of the Omicron variant. Data is shown as median with IQR. **B**, Titers of IgG binding to the RBD protein of the Wuhan strain. Data is shown as mean±SD. **C**, Titers of neutralizing antibodies against the pseudo-virus of the Omicron variant. Data is shown as mean±SD. **D**, Titers of neutralizing antibodies against the pseudo-virus of the Delta variant. Data is shown as mean±SD. Parametric t-test was used to analyze the data in **B**. Non-parametric t-test was used to analyze the data in **A, C** and **D**.

### The S_Omicron DNA vaccine booster improved T cell responses against both the Wuhan strain and the Omicron variant

At the 4^th^ week post the booster vaccinations, mouse splenocytes were freshly isolated and stimulated with peptides covering the full-length spike protein of Wuhan strain or the fragments of Omicron spike protein containing mutations. Specific T cell responses were assessed by intracellular cytokine staining assays (Figure 3 and 4). Our data showed that a booster vaccination of the S_Wuhan DNA vaccine failed to improve specific CD4^+^ T cell responses recognizing spike proteins of both the Wuhan strain and the Omicron variant (Figure 3A-3C, 4A and 4B). In contrast, the booster vaccination of the S_Omicron DNA vaccine significantly enhanced IL-2^+^TNF-a^+^IFN-γ^+^CD4^+^ T cell responses against the spike proteins of Wuhan strain (*P* =0.0240). In comparison with CD4^+^ T cell responses, specific CD8^+^ T cell responses were significantly improved by the booster doses of both the S_Wuhan and the S_Omicron DNA vaccines compared with the control group (Figure 3D-3F, 4C and 4D). It is worth noting that the booster shot of the S_Omicron DNA vaccine elicited the highest IFN-γ^+^CD8^+^, TNF-a^+^CD8^+^, IL-2^-^TNF-a^-^IFN-γ^+^CD8^+^ and IL-2^-^TNF-a^+^IFN-γ^+^CD8^+^ T cell responses against spike proteins of both the ancestral and the Omicron variants among the three groups. However, no significant differences of CD8^+^ T cell responses were observed between the groups boosted with the S_Omicron and the S_Wuhan DNA vaccines.

**Figure 3.**
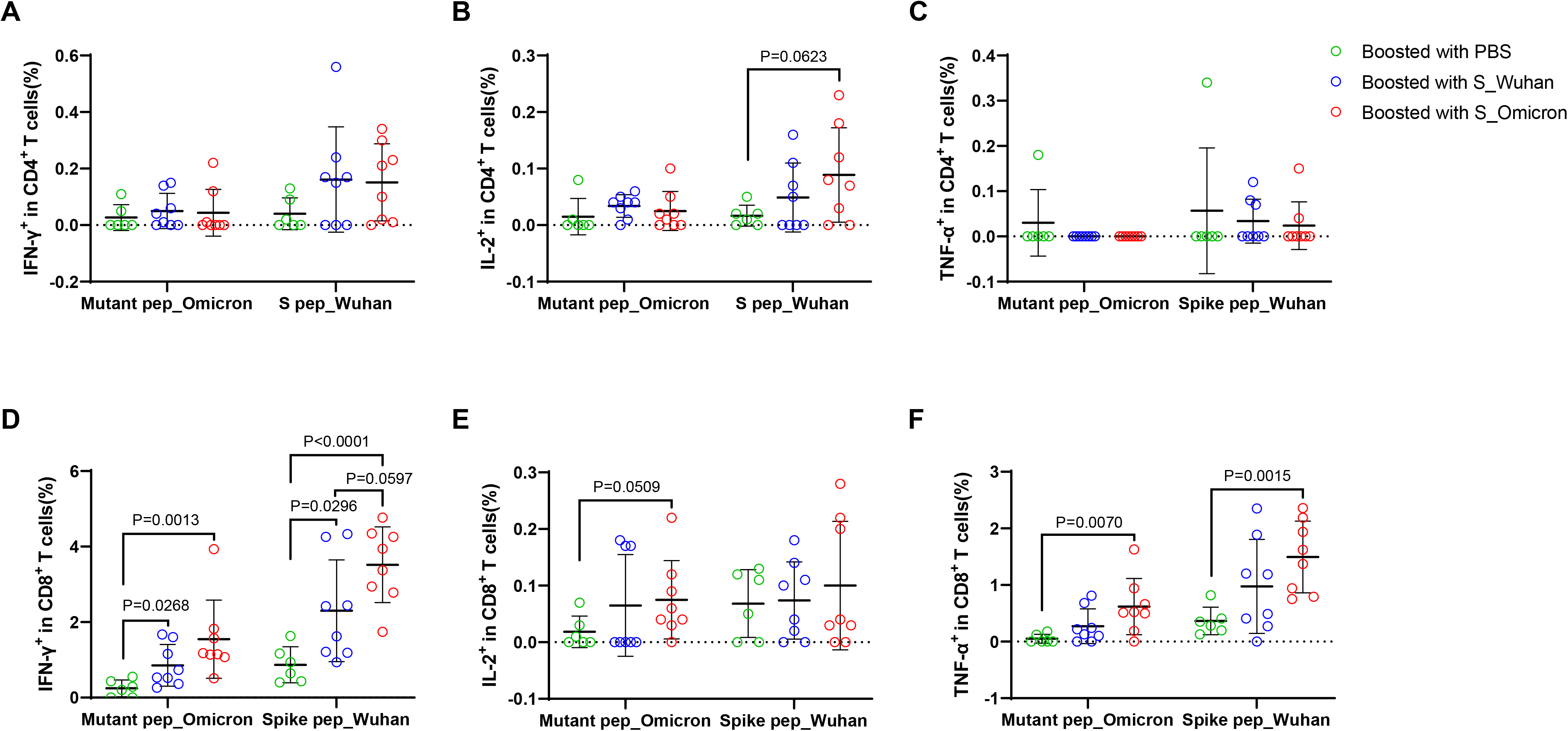
Boosting with the S_Omicron DNA vaccine enhanced the magnitudes of cross-reactive T cell responses. Comparisons of IFN-γ^+^CD4^+^, IL-2^+^CD4^+^ and TNF-a^+^CD4^+^ T cell responses were in **A, B** and **C**, respectively. Comparisons of IFN-γ^+^CD8^+^, IL-2^+^CD8^+^ and TNF-a^+^CD8^+^ T cell responses were in **D, E** and **F**, respectively. Data are shown as mean±SD. Statistical analyses were performed using the method of parametric t-test.

**Figure 4.**
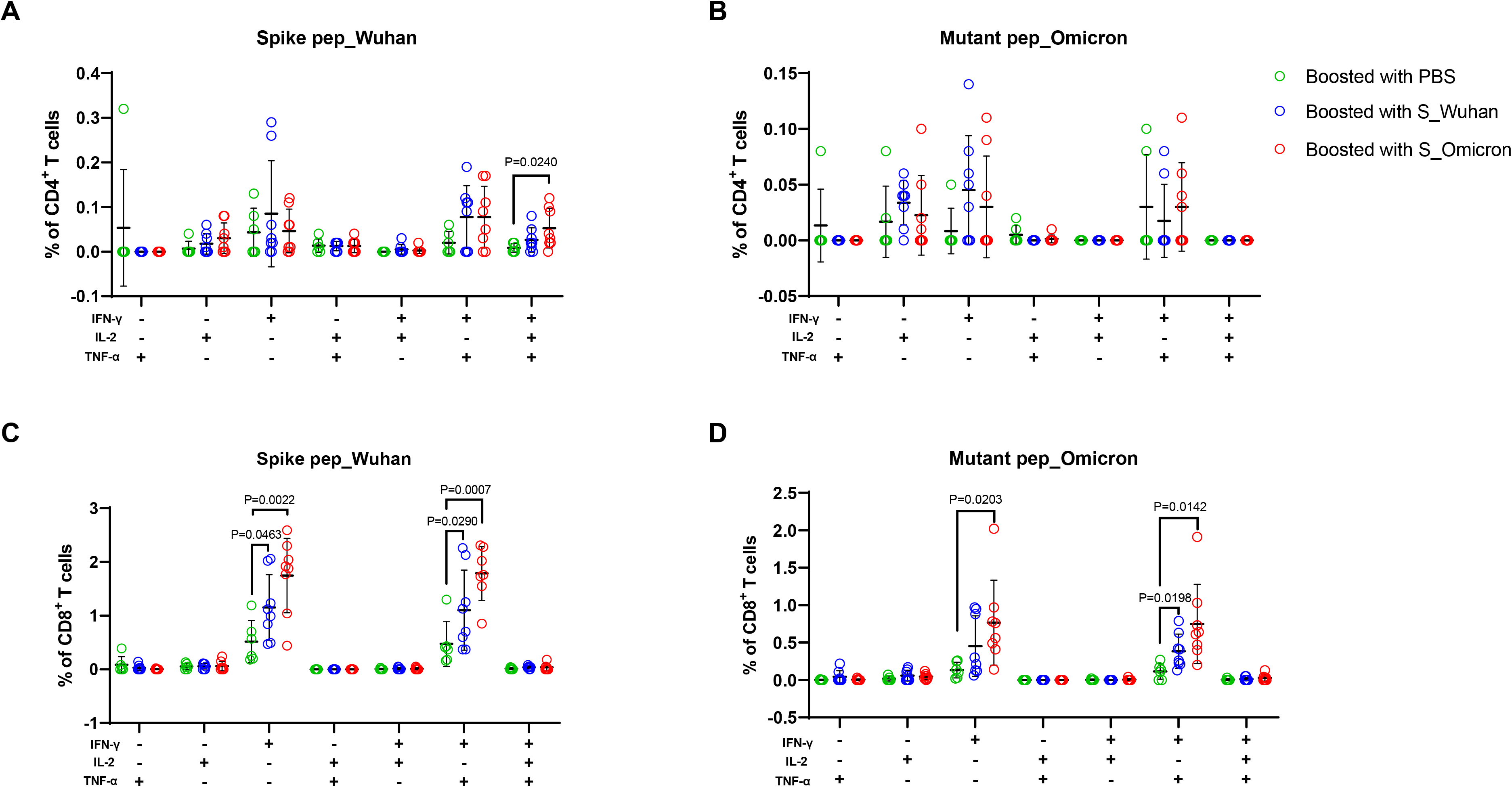
Boosting with the S_Omicron DNA vaccine improved the functionality of spike protein specific T cells. The functionality of spike protein specific T cells was delineated via measuring the secretions of IL-2, TNF-a and IFN-γ. Analyses of the functionality of the Wuhan strain and the Omicron variant specific CD4^+^ T cells were shown in **A** and **B**, respectively. Analyses of the functionality of Wuhan and Omicron specific CD8^+^ T cells were shown in **C** and **D**, respectively. Data are shown as mean±SD. Statistical analyses were performed using the method of parametric t-test.

## Discussion

Extensive mutations in the spike protein render the Omicron variant more likely to escape current vaccines than ancestral SARS-CoV-2 variants (32, 33). A third dose can reduce the risk of hospitalization due to Omicron, however, the effectiveness may diminish quickly (34). The necessity and benefit of an Omicron vaccine is now under debate. Comparisons between current vaccines and Omicron-matched vaccines can provide valuable information on development of new COVID-19 vaccines.

Given that a massive population have been previously infected by and/or vaccinated against SARS-CoV-2 globally, the selection of a proper booster vaccine is critical for eliciting optimal protective immunities against mutated variants. Hence, in this study, we compared the booster effect of a Wuhan S DNA vaccine and an Omicron S DNA vaccine in mice primed with two-dose of a Wuhan S DNA vaccine. In consistent with a previous study (25), our results show that a booster with Omicron-matched vaccine can elicit stronger inhibitory antibody responses against the Omicron variant. Meanwhile, we found that the mice boosted with the Omicron S DNA vaccine also maintained relatively higher RBD binding antibody and neutralizing antibody responses against the Delta variant, suggesting that the Omicron-matched vaccine might help to boost broadly neutralizing antibody responses.

In addition to humoral responses, cross-reactive T cell responses have also been observed in previously infected or vaccinated people (19, 35-37), which hold the potential to protect against severe disease with Omicron. Here we show that the Omicron-matched vaccine is more efficient in boosting cross-reactive T cell responses. As a recent study demonstrates that the Omicron variant might escape T cell immunity in some individuals with prior infection or vaccination (38), a booster vaccination that can improve T cell cross-reactivity will very likely help to protect against emerging SARS-CoV-2 variants. We note two major limitations in our study. First, we were not able to conduct a live-virus challenge experiment, therefore, we do not know whether the observed improvements of humoral and cellular immunities can be translated into superior protection. Second, the results were generated using a mouse model, which might not completely mimic the characteristics of human immune responses. The Omicron variant challenge studies in animals and vaccination in humans will be required for corroboration.

## Author contributions

Y.M.W., C.Q. and W.H.Z. designed the study. L.Q.J., Y.Z, S.S.L, Y.F.Z. and Y.M.W. conducted the experiments. Y.M.W., L.Q.J. and Y.Z. analyzed the data and drafted the manuscript. C.Q., W.H.Z., D.M.Y. and W.H.W. revised the manuscript.

## Acknowledgements

This work was funded by the National Natural Science Foundation of China (Grant No. 81971559, 31872744 and 82041010) and the Science and Technology Commission of Shanghai Municipality (Grant No. 21NL2600100).

## Figure legends

**Supplementary figure 1 Gating strategy of flow cytometry**. Briefly, single cells were identified from the total events via an FSC-H vs FSC-A gating. Next, a gating of FSC vs SSC was used to find the lymphocyte population. Then, CD3^+^ T cells were found via a gating of SSC vs CD3-PE-Cy7, which were further divided into CD3^+^CD4^+^ and CD3^+^CD8^+^ T cells via gating on CD4 vs CD8. Finally, frequencies of specific T cells were measured via gating on IL-2, TNF-a and IFN-γ, respectively.

