## Supplementary figures and images for "Omicron booster in ancestral strain vaccinated mice augments protective immunities against both the Delta and Omicron variants"

### Supplementary figure 1

#### Supplementary figure 1

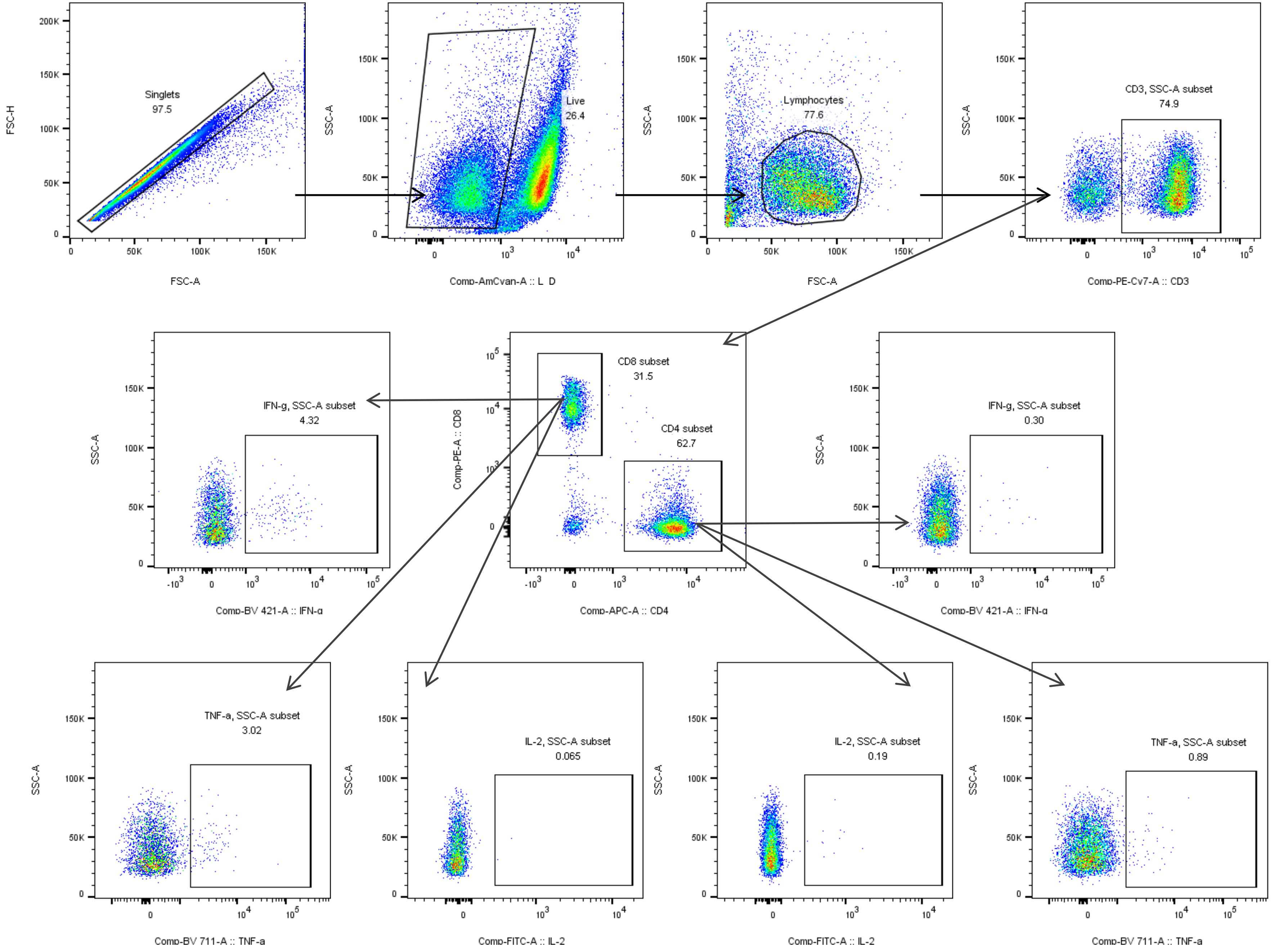
